# Longitudinal data reveal strong genetic and weak non-genetic components of ethnicity-dependent blood DNA methylation levels

**DOI:** 10.1101/339770

**Authors:** Chris McKennan, Katherine Naughton, Catherine Stanhope, Meyer Kattan, George T. O’Connor, Megan T. Sandel, Cynthia M. Visness, Robert A. Wood, Leonard B. Bacharier, Avraham Beigelman, Stephanie Lovinsky-Desir, Alkis Togias, James E. Gern, Dan Nicolae, Carole Ober

## Abstract

Epigenetic architecture is influenced by genetic and environmental factors, but little is known about their relative contributions or longitudinal dynamics. Here, we studied DNA methylation (DNAm) at over 750,000 CpG sites in mononuclear blood cells collected at birth and age 7 from 196 children of primarily self-reported Black and Hispanic ethnicities to study race-associated DNAm patterns. We developed a novel Bayesian method for high dimensional longitudinal data and showed that race-associated DNAm patterns at birth and age 7 are nearly identical. Additionally, we estimated that up to 51% of all self-reported race-associated CpGs had race-dependent DNAm levels that were mediated through local genotype and, quite surprisingly, found that genetic factors explained an overwhelming majority of the variation in DNAm levels at other, previously identified, environmentally-associated CpGs. These results not only indicate that race-associated DNAm patterns in blood are present at birth and are primarily genetically, and not environmentally, determined, but also that DNAm in blood cells overall is robust to many environmental exposures during the first 7 years of life.

## Introduction

DNA methylation (DNAm) in the human genome plays a critical in regulating many cellular processes [1, 2], and altered DNAm patterns have been associated with many diseases, including cancer [3], neurological disorders [4, 5] and asthma [6, 7], to name a few. DNAm itself reflects the contributions of genetic variation [8, 9], exposure histories [10–16], and biological factors such as age [17–26], and has therefore been suggested as a mediator of the effect of these factors on disease outcomes [27, 28].

Recently, results from cross-sectional studies have shown that DNAm in blood cells differs across racial and ethnic groups at birth [29, 30] and later in life [31–34], suggesting that it might contribute to race/ethnicity-associated health disparities [30, 31]. Because racial and ethnic group definitions reflect both common genetic ancestries and shared exposure histories [35], it has been postulated that race/ethnicity-associated blood DNAm patterns are an amalgam of genetic and non-genetic components, and understanding the contribution of each can help inform the relative contribution of genetic and socio-cultural diversity to variation in DNAm levels [31]. For example, a previous study [31] partitioned variation in DNAm levels into genetic and non-genetic sources, and concluded that non-genetic, socio-cultural sources had a significant impact on blood DNAm levels. However, that study, and all previous studies that identified race/ethnicity-associated DNAm marks, relied on cross-sectional data and were therefore not able to asses the temporal stability of those marks. Understanding the stability of race/ethnicity-dependent DNAm present at young ages can help to determine the extent to which race/ethnicity-dependent properties of epigenetic-driven diseases can be attributed to the innate or acquired methylome [29], and identify CpGs whose DNAm is robust or sensitive to accumulated exposures. We therefore sought to fill this gap by first identifying the factors contributing to and the temporal stability of race/ethnicity-dependent blood DNAm levels, and consequently, determining the relative contributions of genetic and environmental factors to the variation in blood DNAm levels in general.

To do so, we studied global DNAm patterns at over 750,000 CpG sites on the Illumina EPIC array in cord blood mononuclear cells (CBMCs) collected at birth and in peripheral blood mononuclear cells (PBMCs) collected at 7 years of age from 196 children participating in the Urban Environment and Childhood Asthma (URECA) birth cohort study [36, 37]. This cohort is part of the NIAID-funded Inner City Asthma Consortium and is comprised of children primarily of Black and Hispanic self-reported ethnicity, with a mother and/or father with a history of at least one allergic disease, and living in low socioeconomic urban areas (see O’Connor et al. [37] for details of enrollment criteria). Mothers of children in the URECA study were enrolled during pregnancy and children were followed from birth through at least 7 years of age.

The longitudinal design of the URECA study provided us with the resolution to partition genetic from non-genetic effects on race/ethnicity-associated DNAm patterns, and yielded new insight into the factors affecting DNAm patterns at CpG sites in mononuclear (immune) cells during early life in ethnically admixed children. Using a novel statistical method that provides a general framework for analyzing longitudinal genetic and epigenetic data, we show that while DNAm levels vary with chronological age, race/ethnicity-dependent DNAm patterns are overwhelmingly conserved over the first 7 years of life and that these patterns are strongly associated, and often mediated, by local genotype. Relatedly, the variation in DNAm levels at previously reported robust exposure-associated CpGs was overwhelmingly dominated by genetic rather than environmental factors in these children. Considering the results of our study and those of a recently published comprehensive review on environmental epigenetics research [38], we suggest that race/ethnicity-dependent blood DNAm levels in particular, and blood DNAm levels in general, are primarily driven by genetic factors, and are not as responsive to environmental exposures as previously suggested [31], at least during the first 7 years of life.

## Results

Our study included 196 children participants in the URECA cohort who had high quality DNA from both CBMCs and PBMCs collected at birth and age 7, respectively, available for our study [36] (see Methods). The URECA children were classified by parent- or guardian-reported race into one of the following categories: Black, *n* = 147; Hispanic, *n* = 39; White, *n* = 1; Mixed race *n* = 7, and Other, *n* = 2. A description of the study population is shown in Table 1. Genetic ancestry, assessed using principle component analysis (PCA), revealed varying proportions of African and European ancestry along PC1 (Figure 1). Because there was little separation along PC2, and no genome-wide significant correlation between PC2 through PC10 and DNAm levels at either age, we defined PC1 as inferred genetic ancestry. The reported races of the children are also shown in Figure 1. We included only the 186 self-reported Black and Hispanic children in subsequent analyses of reported race.

**Table 1:** *Covariates for the n* = 196 *URECA children in our study, stratified by self-reported race.*

|  | Black | Hispanic | White | Mixed | Other |
| --- | --- | --- | --- | --- | --- |
| Sample Size | 147 | 39 | 1 | 7 | 2 |
| Males (%) | 71 (48%) | 25 (64%) | 0 (0%) | 4 (57%) | 0 (0%) |
| Asthma diagnosis at age 7 (%) | 38 (26%) | 12 (31%) | 0 (0%) | 2 (29%) | 0 (0%) |
| Gestational age at birth, in weeks (mean [range]) | 39.0 [34,42] | 38.9 [35,41] | 36.0 | 39.1 [37,40] | 39.0 [38,40] |
| **Sample Collection Site** |  |  |  |  |  |
| Baltimore (%) | 64 (44%) | 1 (3%) | 1 (100%) | 3 (43%) | 2 (100%) |
| Boston (%) | 17 (12%) | 5 (13%) | 0 (0%) | 2 (29%) | 0 (0%) |
| New York (%) | 23 (16%) | 32 (82%) | 0 (0%) | 1 (14%) | 0 (0%) |
| St. Louis (%) | 43 (29%) | 1 (3%) | 0 (0%) | 1 (14%) | 0 (0%) |

**Figure 1:**
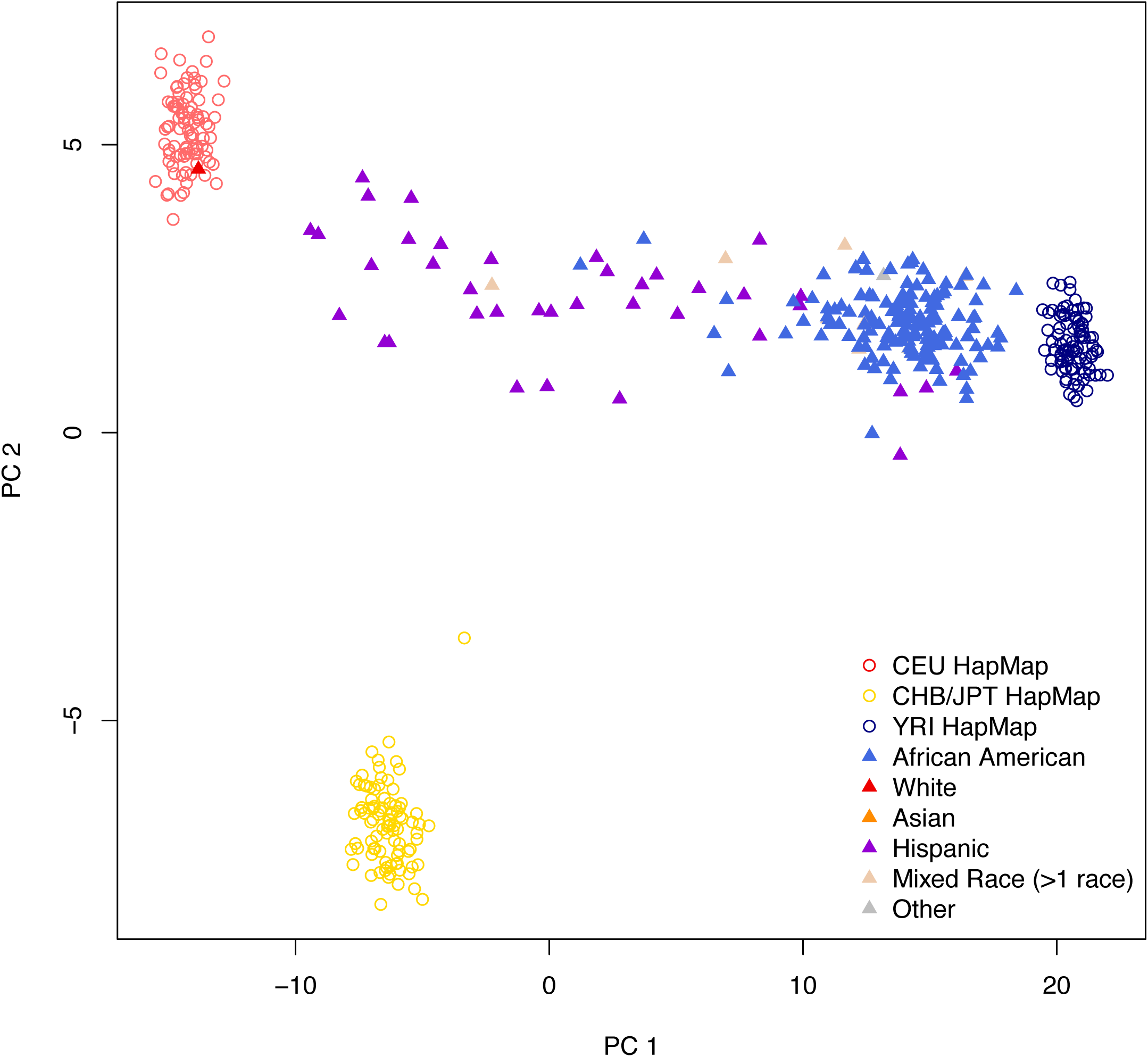
Estimated ancestry principal components (PCs) 1 and 2. Nearly all the variation in ancestry separates along PC1 in the URECA sample. Filled triangles represent the 196 URECA children in this study, with their self-reported race shown in different colors. Open circles are reference control samples from HapMap; red = Utah residents from northern and western Europe (CEU); yellow = east Asian (Chinese and Japanese); dark blue = Africans from Nigeria (Yoruban).

### Reported race effects on DNA methylation patterns are conserved in magnitude and direction between birth and age 7

We first attempted to determine the temporal stability of reported race-associated DNAm patterns by addressing three questions. What is the correlation between reported race and DNAm levels at individual CpG sites at birth and age 7? Is the direction and magnitude of the correlation between reported race and DNAm levels conserved between birth and age 7? Does the correlation between DNAm levels and reported race differ significantly between birth and age 7? While these questions are important in their own right, their answers can also help determine the nature of these reported race-associated patterns. For example, race-associated DNAm levels that differ at birth and age 7 might reflect race-dependent exposure histories, while race-associated DNAm patterns that are conserved may be genetic in nature, since genetically-dependent DNAm patterns are conserved from birth to later childhood [39].

Standard hypothesis testing can be used to answer the first question but is not appropriate for answering the second or third because failure to reject the null hypothesis that the effects are equal at birth and age 7 does not imply the null hypothesis is true. Additionally, because our studies were conducted in CBMCs at birth and PBMCs at age 7, DNAm levels at birth and age 7 may differ slightly due to differences in cell composition [40]. To address these issues, we built a Bayesian model (see Model (1) in Methods) and let the data determine both the strength of the correlation between reported race (based on self-report) and DNAm levels, and how similar the correlations are at birth and age 7. We then answered the above three questions by defining and estimating the conserved (con) and discordant (dis) sign rates for each CpG *g* = 1, …, 784, 484:

> *con*_*g*_ = Posterior probability that CpG *g*’s ancestry effects at birth and age 7 were non-zero, had the same sign AND the sign was estimated correctly.
>
> *dis*_*g*_ = Posterior probability that the ancestry effect for CpG *g* was non-zero at one age and zero or in the opposite direction at the other age.

For a given posterior probability threshold, these quantities partition the ancestry-associated CpGs into two groups: those whose ancestry effects were non-zero and conserved from birth to age 7 and those whose ancestry effects were different at birth and age 7. Detailed descriptions of our model and estimation procedure are provided in the “Joint modeling of DNA methylation at birth and age 7” section in Methods. Supplemental Figure S1 provides insight into how the conserved sign rate compares with standard univariate *P* values.

After fitting the relevant parameters in the model to the data, we were able to estimate the fraction of CpGs with non-zero reported race effects at both ages and assign them into one of four possible bins: the two effects were completely unrelated (ρ = 0), moderately similar (ρ = 1/3), very similar (ρ = 2/3), or identical (ρ = 1). Note that if a non-trivial fraction of CpG sites had ancestry effects that were in opposite directions at birth and age 7, they would be assigned to the first bin (ρ = 0). In fact, we estimated that only 0.2% of the CpGs with non-zero reported effects at both ages had unrelated or moderately similar reported race effects, whereas 30.7% fell in the very similar bin and 69.1% had identical reported race effects at birth and age 7 (Supplemental Figure S2). These data indicate that when reported race effects on DNAm levels are present (i.e., non-zero) at both birth and age 7, they tend to be very similar or exactly the same at both ages with respect to both direction and magnitude.

We then estimated the conserved and discordant sign rates for all 784,484 probes and classified a CpG as a reported race-associated CpG (RR-CpG) if its conserved or discordant sign rate was above 0.80 (i.e. *con*_*g*_ ≥ 0.8 or *dis*_*g*_ ≥ 0.8). At this threshold, we identified 2,162 RR-CpGs, 2,157 (99.8%) of which were conserved in sign (*con*_*g*_ ≥ 0.8). Compared to self-reported Hispanic children, self-reported black children tended to have higher DNAm levels at 1,288 (60%) of the conserved RR-CpGs (*P* = 8.6 × 10^−38^). This trend replicated when we substituted inferred genetic ancestry for reported race and is in accordance with previous observations [6, 33], indicating individuals with more African ancestry tend to have overall more DNAm. Interestingly, there was an under enrichment of RR-CpGs in CpG islands (*P* = 3.10 × 10^−12^), which mirrors the observation that CpGs whose DNAm is under genetic control typically lie outside of CpG islands [41]. The fact that only 5 of the 2,162 RR-CpGs had discordant reported race effects at birth and age 7 (*dis*_*g*_ ≥ 0.8) corroborates the observations made in the previous paragraph and answers the second question in the affirmative: if DNAm levels are correlated with reported race at birth, the magnitude and direction of the correlation is almost certainly conserved at age 7 (and vice-versa).

### Inferred genetic ancestry is more correlated with DNA methylation than is self-reported race

The observed correlations between ancestry and DNAm levels may reflect differences in environmental exposures [31, 33], due to associations between race or ethnicity with socio-cultural, nutritional, and geographic exposures, among others [42]. In fact, a previous cross sectional study suggested that self-reported ethnicity explained a substantial proportion of the variance of blood DNAm levels measured in Latino children of diverse ethnicities [31]. They concluded that ethnicity captured genetic, as well as the socio-cultural and environmental differences, that influence DNAm levels. If this were the case in the URECA children, the effect of inferred genetic ancestry on DNAm levels should be comparable to that of reported race. To assess this possibility in the URECA children, we repeated the analyses described above but substituted inferred genetic ancestry for reported race. This analysis revealed 8,597 inferred genetic ancestry-associated CpGs (IGA-CpGs), of which 8,579 (99.8%) were conserved in sign (*con*_*g*_ ≥ 0.8). This was significantly more than the 2,162 RR-CpGs identified in the reported race analysis above (Figures 2a-b).

**Figure 2:**
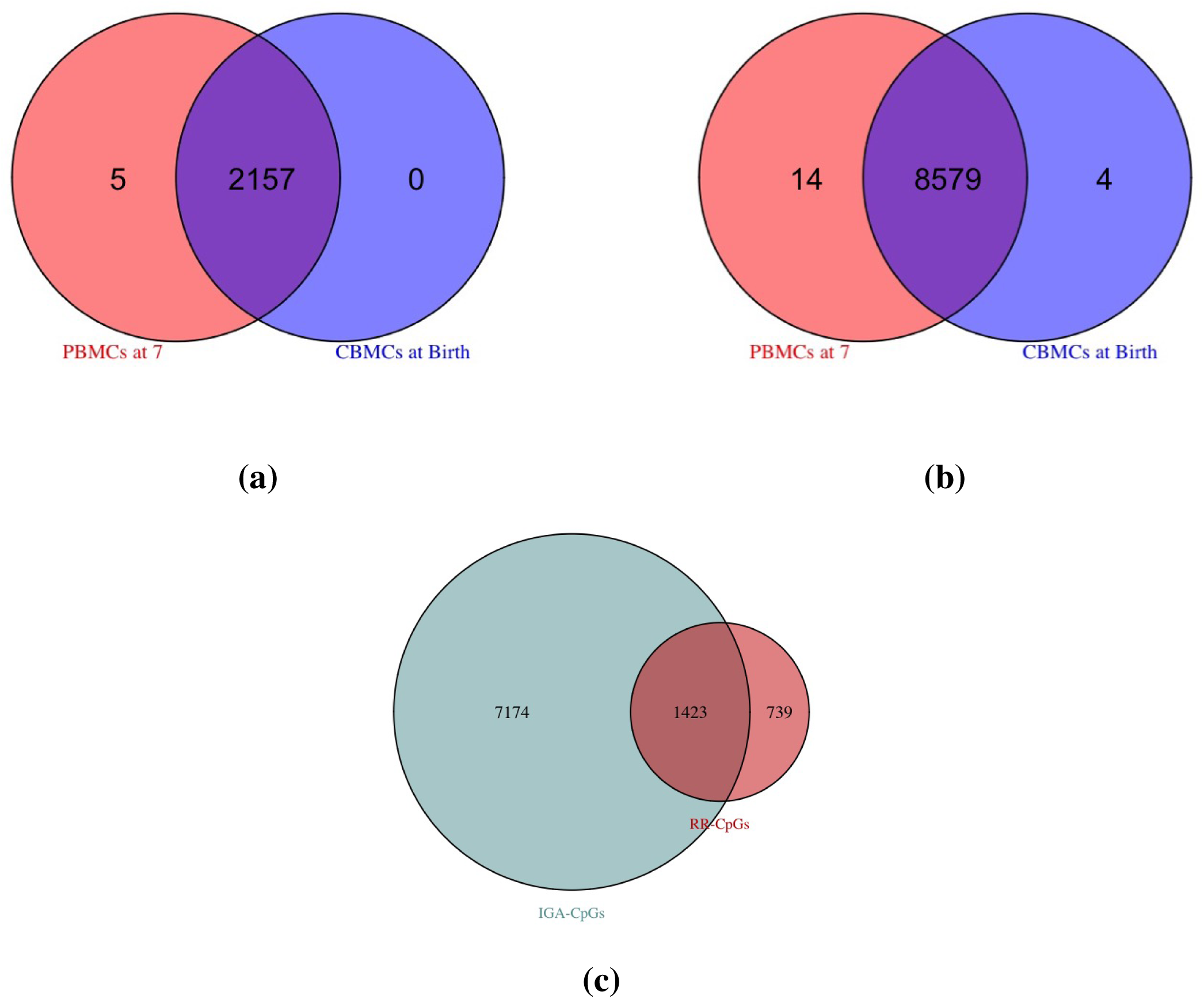
Overlapping ancestry CpGs at birth and at age 7. (a): self-reported race-associated CpGs (RR-CpGs) with *con*_*g*_ ≥ 0.8 (violet) or *dis*_*g*_ ≥ 0.8 (red or blue). A discordant RR-CpG was classified as significant at birth but not at age 7 (blue) if the marginal posterior probability that the effect was non-zero at birth was greater than that at age 7. Discordant RR-CpGs that were significant at age 7 but not at birth (red) were defined analogously. (b): The same as (a), but for inferred genetic ancestry-associated CpGs (IGA-CpGs). (c): The overlap between RR-CpGs (*con*_*g*_ ≥ 0.8 or *dis*_*g*_ ≥ 0.8) and IGA-CpGs (*con*_*g*_ ≥ 0.8 or *dis*_*g*_ ≥ 0.8).

To further explore this finding, we examined the overlap between RR-CpGs and IGA-CpGs (Figure 2c). Because reported race is an estimate of inferred genetic ancestry, there is a substantial overlap between IGA-CpGs and RR-CpGs. Contrary to the results from the previous study [31], which estimated that only 35% of their ethnicity-associated were also genetic ancestry-associated CpGs (Figure 5A in [31]), 66% of RR-CpGs in our study were also IGA-CpGs, and therefore represent only a subset of the IGA-CpGs. This indicates that while IGA-CpGs include most RR-CpGs, reported race does not capture most of the variation in DNAm levels attributable to genetic ancestry in these children.

The differences between our results and those reported in the aforementioned study may be due to the fact that sample collection site explained 80% of the variance in Mexican versus Puerto Rican ethnicity in [31], but was not accounted for in their analyses. The fact that sample collection site was associated with the DNAm levels of 865 CpGs at birth or age 7 at a 5% FDR in our study suggests that sample collection site could have confounded the relationship between ethnicity and DNAm in the previous study (see page 3 in the Supplement for details).

### The association between DNA methylation and reported race is largely genetically driven

To further address the question of whether reported race effects on DNAm levels at either birth or age 7 were primarily due to genetic variation or to environmental exposures, we used local genetic variation (within 5kb of a CpG site) and DNAm data at birth and age 7 in the 147 self-reported Black children in our study to map methylation quantititave trait loci (meQTLs). Of the 519,696 CpGs within 5kb of a SNP, 65,068 and 70,898 had at least one meQTL in CBMCs at birth and in PBMCs at age 7, respectively, at an FDR of 5%. In addition, 51% of all RR-CpGs with at least one SNP in the ±5kb window had at least one meQTL at birth or age 7 at an FDR of 5%, which was a significant enrichment when compared to the 17% observed for non-RR-CpGs (Figure 3a-b).

**Figure 3:**
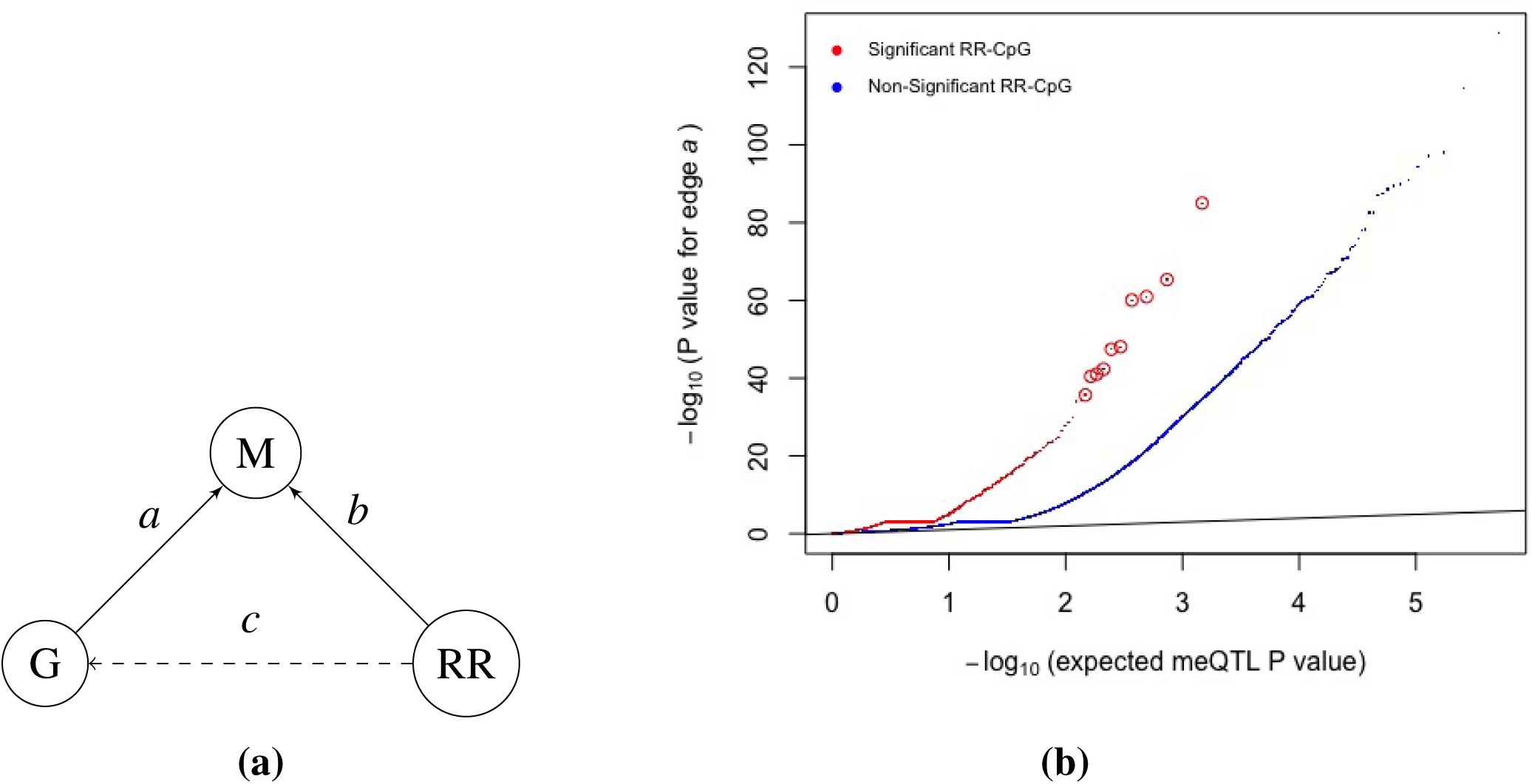
RR-CpGs are enriched for CpGs with meQTLs. (a) Illustration of the causal relationship between the DNAm (M) at a CpG site, the genotype (G) at the SNP within ±5kb of the CpG that had the smallest meQTL *P* value and self-reported race (RR). Each graph corresponds to a unique CpG. (b) Plots of the meQTL *P* value for edge *a* in CBMCs at birth, where CpGs were stratified by whether or not it was an RR-CpG (*con*_*g*_ ≥ 0.8 or *dis*_*g*_ ≥ 0.8). The ten enlarged red circles are just for visual aid.

To provide additional evidence that local genotype mediates the effect of reported race on DNAm levels, we used logistic regression to regress the genotype of each SNP within ±5kb of a RR-CpGs. The goal was to determine the fraction of RR-CpGs at which the observed variation was mediated through local genotype, i.e. RR-CpGs with both edges *a* and *c* in Figure 3a. Since genotype is highly correlated with race, most SNPs will possess edge *c*. Therefore, a reasonable upper bound for this quantity is 51%, the fraction of RR-CpGs with at least one meQTL in their ±5kb window. To determine a lower bound, we used the results of the abovementioned logistic regression to conservatively estimate that at least 26% of all RR-CpGs with at least one SNP in their ±5kb windows had both edges *a* and *c* (see pages 3-5 in the Supplementary Material for calculation details). Interestingly, substituting inferred genetic ancestry for self-reported race in the above analysis yielded nearly identical upper and lower bounds, providing evidence for local genotype mediating the effects of reported race on DNAm levels at RR-CpGs.

### Genetic and biological factors explain most of the variation in blood DNA methylation levels

Given the suggested genetic nature of race/ethnicity-dependent blood cell DNAm levels, we next sought to determine the relative contributions of genetic variation, age and environmental factors on CMBC and PBMC DNAm levels in general at birth and age 7 in the URECA cohort. First, we identified 2,836 gestational age-related CpGs at birth and 16,172 age-related CpGs (CpGs whose DNAm levels changed from birth to age 7) at 5% FDRs. These two sets of CpGs were strongly enriched for CpGs used to predict gestational age in Knight et al. [21] and to predict chronological age in Horvath [18], as well as for CpGs whose blood DNAm levels changed from birth to age 5 in Pérez et al. [43] (see Supplemental Figure S3). Moreover, the estimates of the age effects among age-related CpGs in our study showed the same direction of change as their corresponding estimated gestational age effects at birth in 97% of the 16,172 age-related CpGs. This included 14,186 gestational age-associated effects that were not significant at a 5% FDR threshold but showed the same direction of change. This concordance in direction of effect is unlikely to occur by chance (*P* value < 10^−119^, pages 5-7 of the Supplementary Material for calculation details). Taken together with the enrichments for age-associated CpGs described above, we suggest that the majority of the changes in DNAm levels from birth to age 7 is due to aging-related mechanisms rather than age-dependent environmental exposures.

We next attempted to determine the relative contributions of genetic and environmental factors on DNAm levels in blood. With the exception of maternal cotinine levels during pregnancy, which previously showed robust and reproducible associations with blood DNAm levels at birth [11–15] and in early childhood [10, 13, 16], none of the direct or indirect measures of exposures that were available in this cohort were associated with DNAm levels at either age after adjusting for multiple testing (see pages 1-2 in the Supplementary Material for a complete list). Therefore, in order to maximize our chances of identifying environmental variation in these data, we restricted our analyses to the 6,073 maternal smoking-related CpGs identified in Joubert et al. [15], who performed a meta analysis of maternal smoking during pregnancy on 6,685 infants from 13 cohorts. In our data, DNAm levels at birth and age 7 at 505 (9.2%) and 407 (7.4%) of the 5,500 maternal smoking-related CpGs that passed QC in our study, respectively, were nominally correlated (*P* value ≤ 0.05) with maternal cotinine levels (enrichment *P* values = 7.08 × 10^−34^ and 6.49 × 10^−8^). While this enrichment was not unexpected, we were surprised to observe that the maternal smoking-related CpGs were enriched for meQTLs (Figure 4a). Additionally, there was a strong enrichment of the 8,579 conserved inferred genetic ancestry-associated CpGs among the 5,500 maternal smoking-related CpGs that passed QC in our study (fold enrichment = 2.53; *P* value = 6.42 × 10^−33^), indicating the maternal smoking-related CpGs were enriched for genetically regulated CpGs. Furthermore, genotype at the closest SNP for over 95% of the maternal smoking-related CpGs explained a greater proportion of the variance in DNAm levels at birth than did maternal cotinine levels (Figure 4b, see pages 7-9 in the Supplementary Material for analysis details). These results were identical for DNAm measured at age 7, and showed that genetic, and not environmental, factors are responsible for the majority of the variation in DNAm levels at even the most robust and replicated environmentally-associated CpGs in these children.

**Figure 4:**
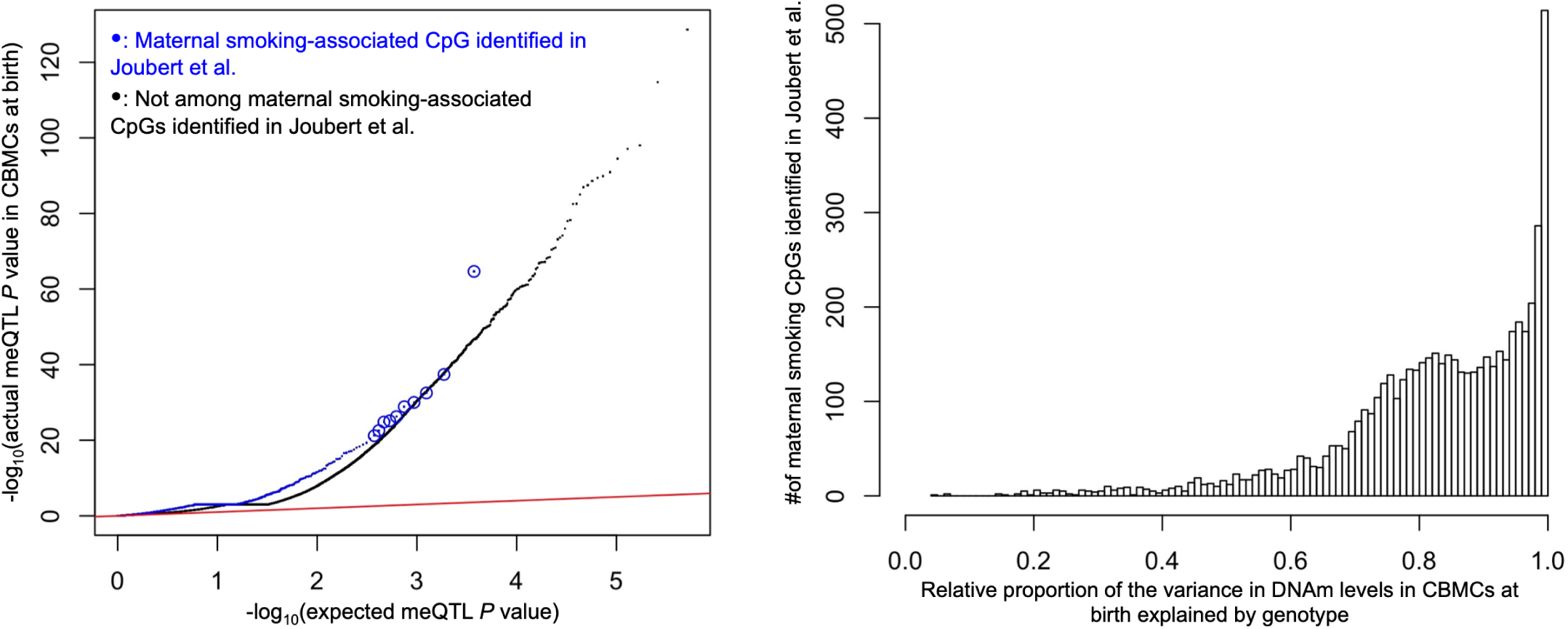
meQTL *P* value enrichment, where circled blue points are for visual aid (left), and the relative proportion of variance in DNAm levels explained by genotype (right). The x-axis of the latter was defined as the ratio of the proportion of variance in DNAm levels explained by the genotype of each CpG’s closest SNP to the sum of the aforementioned genetic proportion and the proportion explained by maternal cotinine levels during pregnancy. A ratio > 0.5 indicates that local genotype explained more variance than maternal cotinine levels during pregnancy.

## Discussion

The relationships between DNAm, chronological age, and race/ethnicity have the potential to shed light on disease etiology and may help determine the relative genetic and environmental contributions to the observed inter-individual variability of the epigenome [17–23, 29–34]. While it has previously been shown that race/ethnicity is related to DNAm in cross-sectional studies [29–34] and that statistically significant meQTLs are conserved as individuals age [39], it has yet to be shown that race/ethnicity-dependent DNAm marks are conserved as children age, and relatedly, that exposure histories explain a comparatively small fraction of the variation in DNAm levels.

Even though there was substantial change in blood DNAm levels over time among children in this cohort, self-reported race effects on DNAm were overwhelmingly conserved in both direction and magnitude from birth to age 7. This result, as well as our novel Bayesian inference paradigm used to obtain it, is important in and of itself because it provides an example of, and a general method for identifying, DNAm patterns that are conserved over time, and differentiating between environmentally responsive and temporally stable DNAm marks, which has been highlighted as both a gap in current knowledge and a critical area of future epigenetic research [44]. The consistency of our estimates for inferred genetic ancestry and reported race effects on DNAm levels also demonstrates the fidelity of our processing pipeline that accounts for unobserved factors, including cell composition, because failure to account for latent covariates can lead to biased and irreproducible estimates [45, 46].

While the observation that reported race effects are conserved from birth to age 7 gives credence to the hypothesis that the effects are genetic in nature, it does not rule out the possibility of environmental components or gene-environment interactions that could result in race/ethnicity-associated DNAm patterns prior to birth that persist as the child ages. It was therefore interesting to find that there was a significant under enrichment of RR-CpGs in CpG islands, which agrees with the under enrichment previously observed for CpGs under genetic control [41]. To further explore this, we showed that the RR-CpGs were enriched among CpGs with meQTLs identified in our study, indicating that DNAm levels at many of the RR-CpGs are mediated by local genotype and that much of the reported race-DNAm correlation could be attributed to genetic variation. Moreover, the RR-CpGs were only a small subset of inferred genetic ancestry associated CpGs (IGA-CpGs) in our study. This is contrary to the findings of Galanter et al. [31], who argued that ethnicity-dependent DNAm patterns in admixed populations capture both genetic variation and differences in accumulated exposures. Our results provide evidence for genetics accounting for an overwhelming majority of the correlation between DNAm levels and reported race, which suggests the non-genetic contribution to variability in blood DNAm levels may be smaller than previously thought.

There were several other notable features in these data connoting that genetic, and not environmental, factors were most responsible of the variation in blood DNAm levels in these children. The first was that although average DNAm levels of 16,172 CpGs changed significantly from birth to age 7, the direction of the change in 97% of those CpGs matched the direction of the corresponding correlation between DNAm levels and gestational age at birth. This manifest concordance in the “epigenetic clocks” present at birth and later in life, along with the observation that the 16,172 age-related CpGs were enriched for CpGs used to predict gestational and chronological age, suggests these age-related changes are coordinated by age-related mechanisms, and not due to age-dependent environmental exposures. Second, with the exception of maternal cotinine levels during pregnancy, none of the direct or indirect measures of exposure history were associated with DNAm levels at birth or age 7. This observation is congruent with the results of a recent comprehensive review on environmental epigenetics research, which suggested that the effects of many environmental exposures on DNAm in blood are probably too small to estimate with even large sample sizes [38].

The third, and possibly most surprising, observation in support of strong genetically- and weak environmentally-determined blood DNAm levels was that genetic, and not maternal cotinine levels, were most responsible for the variation in DNAm levels at over 95% of the maternal smoking-associated CpGs identified in Joubert et al. [15]. This is consistent with, and significantly extends, the results in Gonseth et al. [47], which identified genome-wide significant meQTLs for three of the top ten most significant maternal smoking CpGs identified in the Joubert et al. study. One possibility explanation for our observation, as demonstrated in the Gonseth et al. study, is that genotype confounds the relationship between maternal smoking and DNAm. While we did not have sufficient data to confirm this here, it remains an important area of future investigation.

In summary, the results of our study suggest that DNAm levels in blood cells are fairly robust to environmental exposures, including those that are correlated with self-reported race. A better understanding of tissue-specific DNAm responses to environmental exposures could inform the design of future studies and provide insights into the mechanisms through which exposures and gene-environment interactions influence health and disease.

## Materials and methods

### Sample composition

URECA is a birth cohort study initiated in 2005 in Baltimore, Boston, New York City and St. Louis under the NIAID-funded Inner City Asthma Consortium [36]. Pregnant women were recruited. Either they or the father of their unborn child had a history of asthma, allergic rhinitis, or eczema, and deliveries prior to 34 weeks gestation were excluded (see Gern et al. [36] for full entry criteria). Informed consent was obtained from the women at enrollment and from the parent or legal guardian of the infant after birth.

Maternal questionnaires were administered prenatally and child health questionnaires administered to a parent or caregiver every 3 months through age 7 years. Gestational age at birth and obstetric history were obtained from medical records. Additional details on study design are described in Gern et al. [36]. Frozen paired cord blood mononuclear cells (CBMCs) and peripheral blood mononuclear cells (PBMCs) at age 7, were available for 196 of the 560 URECA children after completing other studies. After QC, DNAm data were available for 194 children at birth, 195 children at age 7, and 193 children at both time points; genotype data were available in 193 children (194 at birth; 195 at age 7). The sample size for each analysis is given in Table 2.

**Table 2:**
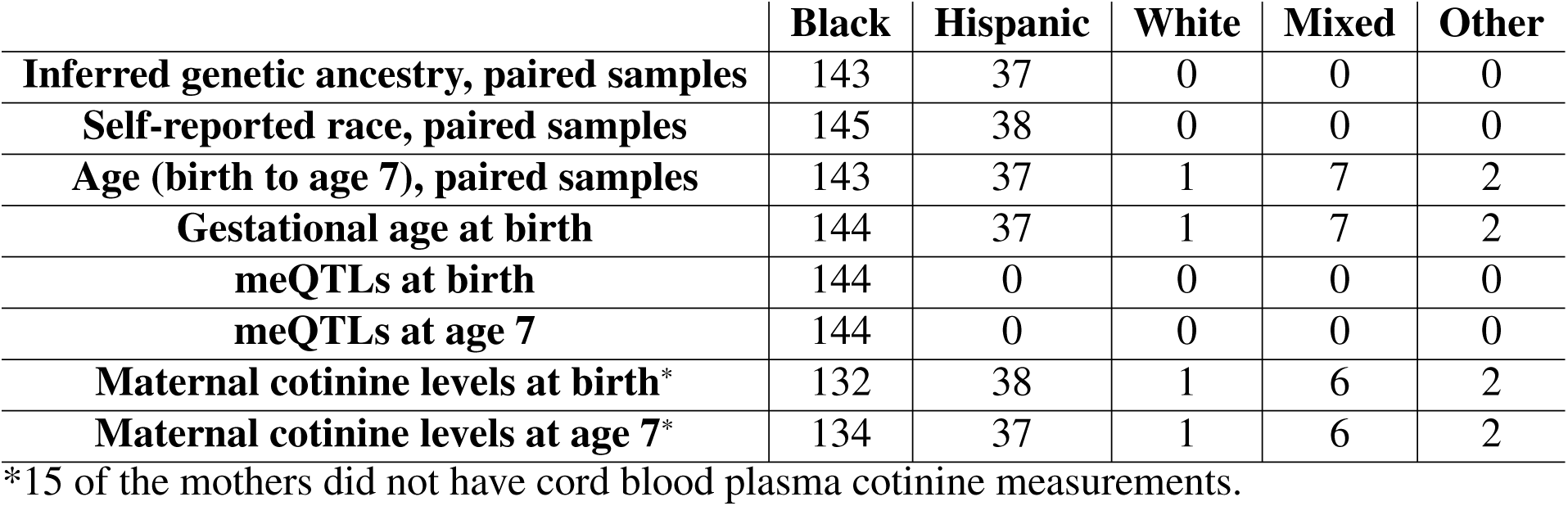
Sample size and composition for each analysis.

|  | Black | Hispanic | White | Mixed | Other |
| --- | --- | --- | --- | --- | --- |
| **Inferred genetic ancestry, paired samples** | 143 | 37 | 0 | 0 | 0 |
| **Self-reported race, paired samples** | 145 | 38 | 0 | 0 | 0 |
| **Age (birth to age 7), paired samples** | 143 | 37 | 1 | 7 | 2 |
| **Gestational age at birth** | 144 | 37 | 1 | 7 | 2 |
| **meQTLs at birth** | 144 | 0 | 0 | 0 | 0 |
| **meQTLs at age 7** | 144 | 0 | 0 | 0 | 0 |
| **Maternal cotinine levels at birth\*** | 132 | 38 | 1 | 6 | 2 |
| **Maternal cotinine levels at age 7\*** | 134 | 37 | 1 | 6 | 2 |

\*15 of the mothers did not have cord blood plasma cotinine measurements.

Maternal cotinine levels were measured in the cord blood plasma at birth, and we categorized mothers as smokers (≥ 10ng/mL; *n* = 31) or non-smokers (< 10ng/mL; *n* = 150), where cotinine levels were missing in 15 mothers. The 10ng/mL threshold was the same as that used in Joubert et al. [15] to define a pregnant mother with a sustained smoking habit, where 147/150 (98%) of the non-smokers in our data had cotinine levels below 2ng/mL, the detection limit of the assay.

### DNA methylation

DNA for methylation studies was extracted from thawed CBMCs and PBMCs using the Qiagen AllPrep kit (QIAGEN, Valencia, CA). Genome-wide DNA methylation was assessed using the Illumina Infinium MethylationEPIC BeadChip (Illumina, San Diego, CA) at the University of Chicago Functional Genomics Facility (UC-FGF). Birth and 7-year samples from the same child were assayed on the same chip and the data were processed using Minfi [48]; Infinium type I and type II probe bias were corrected using SWAN [49]. Raw probe values were corrected for color imbalance and background by control normalization. Three out of the 392 samples (two at birth and one at age 7) were removed as outliers following normalization. We removed 82,352 probes that mapped either to the sex chromosomes or to more than one location in a bisulfite-converted genome, had detection *P* values greater than 0.01% in 25% or more of the samples, or overlapped with known SNPs with minor allele frequency of at least 5% in African, American or European populations. After processing, 784,484 probes were retained and M-values were used for all downstream analyses, which were computed as log2 (methylated intensity +100)− log2 (unmethylated intensity +100). The offset of 100 was recommended in Du et al. [50].

### Genotyping

DNA from the 196 URECA children was genotyped with the Illumina Infinium CoreExome+Custom array. Of the 532,992 autosomal SNPs on the array, 531,755 passed Quality control (QC) (excluding SNPs with call rate < 95%, Hardy-Weinberg *P* values < 10^−5^, and heterozygosity outliers). We conducted all analyses in 293,696 autosomal SNPs with a minor allele frequency ≥ 5%. Genotypes for three children failed QC and were excluded from subsequent analysis that involved genotypes, including methylation quantitative locus (meQTL) mapping, inferred genetic ancestry, or used genetic ancestry PC1 as a covariate. These three children were included in all other analyses.

### Estimating inferred genetic ancestry

Ancestral principal component analysis (PCA) was performed using a set of 801 ancestry informative markers (AIMs) from Tandon et al. [51] that were genotyped in both the URECA children and in HapMap [52] release 23.

### Univariate statistical methods

To determine the effect of gestational age and maternal cotinine levels (smoker vs. non-smokers) on DNAm levels in CBMCs at birth or PBMCs at age 7, we used standard linear regression models with the child’s gender, sample collection site, inferred genetic ancestry and methylation plate number as covariates in our model. We controlled for gestational age in the maternal cotinine analysis. We also estimated cell composition and other unobserved confounding factors using a method described in McKennan et al. [53]. We then computed *P* values for each CpG site and used q-values [54] to control the false discovery rate at a nominal level. We took the same approach to determine CpGs whose DNAm changed from birth to age 7, except the response was measured as the difference in DNAm at birth and age 7. In this analysis, we included the child’s gender, gestational age at birth, inferred genetic ancestry and sample collection site as covariates. Because all paired samples were on the same plate, we did not include plate number as a covariate in this analysis. We also estimated unobserved factors that influence differences in DNAm at birth and age 7 using McKennan et al. [53] and included these latent factors in our linear model.

### Joint modelling of DNA methylation at birth and age 7

We used data from the self-reported Hispanic and Black individuals with DNAm measured at both time points to analyze the effect of ancestry on DNAm levels at CpGs *g* = 1, …, *p* = 784, 484 using the following model:

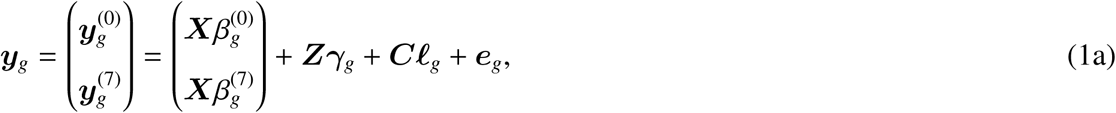

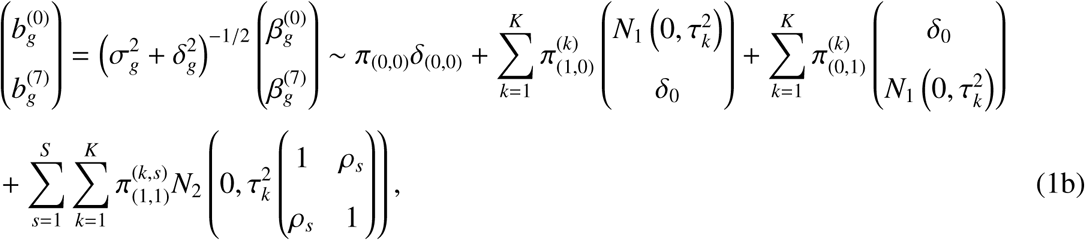

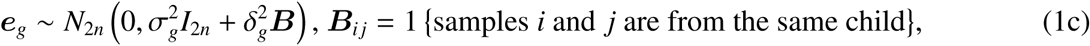

where *δ*_0_ and *δ*_(0,0)_ are the point masses at 0 ∈ ℝ and (0, 0) ∈ ℝ^2^. The vector 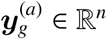 contained the DNAm levels at CpG *g* at age *a*, ***X*** ∈ ℝ^*n*^ contained each child’s inferred genetic ancestry or self-reported race and 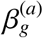 was the effect due to ancestry at age *a*. ***X*** was standardized to have variance 1 when ***X*** was inferred genetic ancestry. The nuisance covariates ***Z*** contained an intercept for the cord blood and PBMC samples, sample collection site, gender, gestational age at birth and plate number. Since gestational age was only correlated with cord blood DNAm, we assumed the effect of gestational age on DNAm at age 7 was zero for all CpG sites. We estimated the unobserved covariates ***C*** with McKennan et al. [55], which accounts for the correlation between samples from the same child.

The entries of the weight vector 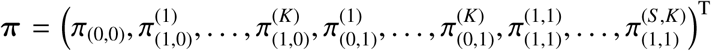 sum to 1, where we set *K* = 5 and *S* = 4. Similar to Flutre et al. [56] and Stephens [57], we specified a grid of correlation coefficients *ρ*_*s*_ ∈ {0, 1/3, 2/3, 1} and a dense grid of effect sizes *τ*_*k*_ ∈ {0.05, 0.1, 0.15, 0.20, 0.25} when ***X*** was inferred genetic ancestry and *τ*_*k*_ ∈ {0.1, 0.15, 0.225, 0.3, 0.375} when ***X*** was reported race. We set *τ*_4_ by first performing a univariate analysis and then estimating the variance of the effect sizes for CpGs with q-values ≤ 0.05, and *τ*_1_ was such that if 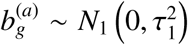, the expected number of CpGs significant at the Bonferroni threshold 0.05/*p* in a univariate analysis would be smaller than 1 for *a* = 0, 7. The proportion of CpGs with non-zero reported race effects at both ages that fell in bin *s* = 1, …, 4 was defined as 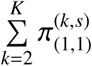, where we ignored the proportion when *k* = 1, because *τ*_1_ was too small to differentiate from zero. The estimated proportion of CpGs in the *ρ*_*s*_ = 2/3 or *ρ*_*s*_ = 1 bins was still over 98% when we included *τ*_1_.

To fit the model, we first regressed out ***Z*** and the estimated ***C*** from both ***y***_*g*_ and ***X*** ⊕ ***X*** and used the residuals in the downstream analysis. We estimated 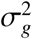 and 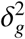 for each *g* = 1, …, *p* with restricted maximum likelihood (REML) and followed Stephens [57] and estimated ***π*** by empirical Bayes via expectation maximization. Supplemental Figures S2 and S4 plot the estimate for ***π*** in the reported race analysis. We then defined *con*_*g*_ and *dis*_*g*_ for each CpG *g* = 1, …, *p* as

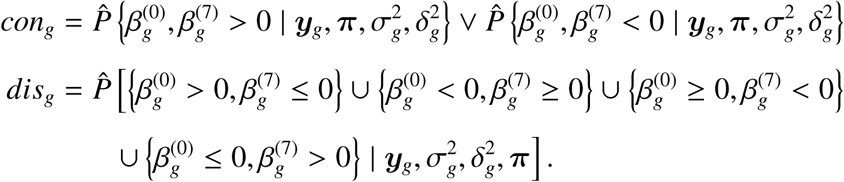

### Determining meQTLs

We performed meQTL mapping in the 145 genotyped, self-reported Black children using the set of 269,622 SNPs with 100% genotype call rate in this subset. We restricted ourselves to this subset of samples to minimize heterogeneity in effect sizes. To identify CpG-SNP pairs, we considered SNPs within 5kb of each CpG, as this region has been previously shown to contain the majority of genetic variability in DNAm [8] and is small enough to mitigate the multiple testing burden, and computed a *P* value for the effect of the genotype at a single SNP on DNAm at the corresponding CpG with ordinary least squares. We then defined the meQTL for each CpG site as the SNP with the lowest *P* value. In addition to genotype, we included inferred genetic ancestry (i.e., ancestry PC1), gestational age at birth, gender, sample collection site and methylation plate number in the linear model, along with the first nine principal components of the residual DNAm data matrix after regressing out the intercept and the five additional covariates. We then tested the null hypothesis that a CpG did not have an meQTL in the 10kb region by using the minimum marginal *P* value in the region as the test statistic and computed its significance via bootstrap. We lastly used q-values to control the false discovery rate.

## Ethical statement

We used de-identified single nucleotide polymorphism, DNA methylation and phenotype data from samples taken from human subjects as part of the Urban Environment and Childhood Asthma study. The WIRB approved human samples to be used in the Urban Environment and Childhood Asthma study (WIRB project number: 20142570).

