## Supplementary material for "Longitudinal data reveal strong genetic and weak non-genetic components of ethnicity-dependent blood DNA methylation levels"

April 2, 2020

This supplement contains a description of additional analyses, additional statistical methods and all supplemental figures referenced in the main text of the manuscript.

### Detailed sample description and additional analyses

#### A complete description of the available covariate information

Maternal questionnaires, including those on smoking, stress, and depression, were administered prenatally and annually after the child's birth. Postnatal child health questionnaires were administered to a parent every three months through age 7 years. Annual visits of child and parent to the study center, starting at one year of age, included questionnaires, anthropomorphic measurements, and phlebotomy. Questionnaires included the Perceived Stress Scale [1], the Edinburgh Postpartum Depression Scale [2], and additional questionnaires to assess stress related to neighborhood factors, violence, and economic hardship [3]. Parent-reported colds were ascertained by telephone questionnaire every three months throughout the first three years of life. Gestational age at birth, maternal infections during pregnancy, and obstetric history were obtained from medical records. Bedroom allergens were measured in vacuumed dust from the child's bedroom, and cotinine levels were measured in cord blood plasma. Pet ownership, number of smokers in the household, daycare attendance, number of siblings, and maternal asthma were ascertained by interview with the mother. Allergic sensitization was determined by prick skin testing for 14 common aeroallergens at 3, 5, or 7 years of age. Aeroallergen sensitization was defined as a wheal $\geq$ 3mm larger than the saline control on prick skin testing or specific IgE $\geq$ 0.35kU/L.

#### Additional analyses to identify exposure-associated CpG sites

We performed additional analyses on every measured direct or indirect measures of environmental exposures to attempt to identify exposures that correlated with DNA methylation (DNAm) levels at either birth or age 7. These exposures were maternal asthma, maternal infections during pregnancy, pet ownership, bedroom allergens, mother stress, anxiety and depression metrics, number of smokers in the household, number of siblings, number of previous live births, daycare attendance, number of colds at age 2 or 3, and allergic sensitization or asthma in the child. Unlike maternal cotinine levels measured during pregnancy, there were no CpGs associated with DNAm levels at birth or age 7 at a 5% FDR.

##### Identifying sample collection site-associated CpG sites

We next attempted to identify CpGs whose DNAm levels at birth or age 7 were associated with sample collection site, which was used to argue that sample collection site might be confounding the relationship between ethnicity and DNAm levels in Galanter et al. [4]. We restricted the analysis to samples from self-reported black and Hispanic children, and regressed methylation at birth or age 7 onto sample collection site (a factor variable with four levels), while accounting for self-reported race (black or Hispanic), sex (male or female), gestational age, inferred genetic ancestry, methylation plate number (a factor variable with five levels) and nine latent factors estimated with the method proposed in McKennan et al. [5]. We identified 865 CpGs whose DNAm levels were significantly correlated with methylation at birth or age 7 at a 5% FDR. To test for differences in the effect due to sample collection site at birth and age 7, we regressed the difference in methylation at birth and age 7 onto sample collection site while accounting for the aforementioned covariates. There were no CpGs at which the effect due to sample collection site differed at birth and age 7 at a 5% FDR.

#### Additional statistical methods

##### A lower bound on fraction of reported race-associated CpGs mediated through local genotype

Here we describe how we conservatively estimated the fraction of reported race-associated CpGs (RR-CpGs) with a SNP in a 10kB window that were mediated by a neighboring SNP. Fix some network composed of a CpG with methylation M and the SNP whose genotype G was most correlated with M according to the meQTL discovery procedure described in Methods. Let {RR → M} be the event the CpG is an RR-CpG, {RR → G} the event RR affects genotype, and {G → M} the event G affects M independently of RR. We would like to estimate

$$\begin{aligned} P(\mathrm{RR} \rightarrow \mathrm{G}, \mathrm{G} \rightarrow \mathrm{M} \mid \mathrm{RR} \rightarrow \mathrm{M}) &= \frac{P(\mathrm{RR} \rightarrow \mathrm{G}, \mathrm{G} \rightarrow \mathrm{M}, \mathrm{RR} \rightarrow \mathrm{M})}{P(\mathrm{RR} \rightarrow \mathrm{M})} \\ &= \frac{P(\mathrm{RR} \rightarrow \mathrm{M} \mid \mathrm{RR} \rightarrow \mathrm{G}, \mathrm{G} \rightarrow \mathrm{M})\, P(\mathrm{RR} \rightarrow \mathrm{G}, \mathrm{G} \rightarrow \mathrm{M})}{P(\mathrm{RR} \rightarrow \mathrm{M})} \end{aligned} \quad \text{(S1)}$$

Define $H_0 = \{\mathrm{RR} \not\rightarrow \mathrm{G}\}$. For each SNP we computed a $P$ value for the null hypothesis $H_0$ using the logistic regression model G ∼ RR, where RR was either Black or Hispanic (we assumed a Hardy-Weinberg equilibrium model for the genotypes of all SNPs considered). Let $t$ be the test statistic from the regression and $t^*_\alpha > 0$ be some threshold with significance level $\alpha$. Then because G and RR are independent under $H_0$ (regardless of whether or not {G → M} or {RR → M} hold),

$$\begin{aligned} q = P(H_0 \mid |t| \geq t^*_\alpha, \mathrm{G} \rightarrow \mathrm{M}, \mathrm{RR} \rightarrow \mathrm{M}) &= \frac{P(|t| \geq t^*_\alpha \mid H_0, \mathrm{G} \rightarrow \mathrm{M}, \mathrm{RR} \rightarrow \mathrm{M})\, P(H_0 \mid \mathrm{G} \rightarrow \mathrm{M}, \mathrm{RR} \rightarrow \mathrm{M})}{P(|t| \geq t^*_\alpha \mid \mathrm{G} \rightarrow \mathrm{M}, \mathrm{RR} \rightarrow \mathrm{M})} \\ &= \frac{P(|t| \geq t^*_\alpha \mid H_0)\, P(H_0 \mid \mathrm{G} \rightarrow \mathrm{M}, \mathrm{RR} \rightarrow \mathrm{M})}{P(|t| \geq t^*_\alpha \mid \mathrm{G} \rightarrow \mathrm{M}, \mathrm{RR} \rightarrow \mathrm{M})} \\ &\leq \frac{\alpha}{P(|t| \geq t^*_\alpha \mid \mathrm{G} \rightarrow \mathrm{M}, \mathrm{RR} \rightarrow \mathrm{M})} \end{aligned}$$

where the equality in the second line comes from the fact that under the null hypothesis and given the rest of the graph, the behavior of G and RR are independent. We therefore upper-bounded $q$ by estimating $P(|t| \geq t^*_\alpha \mid \mathrm{G} \rightarrow \mathrm{M}, \mathrm{RR} \rightarrow \mathrm{M})$ using the RR-meQTL logistic regression test statistics (this is just the Benjamini-Hochberg procedure interpreted in a Bayesian framework). We finally established an estimated lower bound for (S1) by using the following:

$$P(\mathrm{RR} \rightarrow \mathrm{G}, \mathrm{G} \rightarrow \mathrm{M} \mid \mathrm{RR} \rightarrow \mathrm{M}) = \frac{\#\text{ networks with RR} \rightarrow \text{G, G} \rightarrow \text{M, RR} \rightarrow \text{M}}{\#\text{ networks with RR} \rightarrow \text{M}}$$

$$\geq \frac{\#\text{ networks with RR} \rightarrow \text{G, G} \rightarrow \text{M, RR} \rightarrow \text{M and } q \leq 0.2}{\#\text{ of networks with RR} \rightarrow \text{M}}$$
$$\gtrsim (1-0.2)\frac{\#\text{ of networks with } q \leq 0.2 \text{ among RR-meQTLs}}{\#\text{ RR-CpGs}}$$
$$= 0.26.$$

##### Calculating the *P* value for the overlap between gestational age- and chronological age-associated CpGs with the same effect sign

Define $\boldsymbol{y}_g^{(a)}$ to be the DNAm for CpG $g = 1, \dots, 784,484$ at age $a = 0, 7$. We estimated the effect of gestational age $\boldsymbol{X} \in \mathbb{R}^n$ on $\boldsymbol{y}_g^{(0)}$ in the model

$$\boldsymbol{y}_g^{(0)} = \boldsymbol{X}\beta_g^{\text{GA}} + \boldsymbol{Z}_0\boldsymbol{\gamma}_g + \boldsymbol{C}_0\boldsymbol{\ell}_g + \boldsymbol{e}_g^{(0)}, \quad \boldsymbol{e}_g^{(0)} \sim N_n\left(0, \left(\sigma_g^2 + \delta_g^2\right) I_n\right) \quad (g = 1, \dots, p)$$

and the effect of age, $\beta_g^{(0\rightarrow 7)}$, in the model

$$\boldsymbol{y}_g^{(7)} - \boldsymbol{y}_g^{(0)} = \boldsymbol{1}_n\beta_g^{(0\rightarrow 7)} + \boldsymbol{Z}_{\text{diff}}\boldsymbol{\gamma}_g + \boldsymbol{C}_{\text{diff}}\boldsymbol{\ell}_g + \boldsymbol{e}_g^{(\text{diff})}, \quad \boldsymbol{e}_g \sim N_n\left(0, \sigma_g^2 I_n\right) \quad (g = 1, \dots, p)$$

using ordinary least squares, where the nuisance covariates in $\boldsymbol{Z}_0, \boldsymbol{Z}_{\text{diff}}$ are given in Methods and $\boldsymbol{C}_0, \boldsymbol{C}_{\text{diff}}$ were estimated using McKennan et al. [5]. Define the estimated gestational age and age effects to be $\hat{\beta}_g^{\text{GA}}$ and $\hat{\beta}_g^{(0\rightarrow 7)}$, respectively. We use the output of these two regressions to get an approximate upper bound for the expected number of pairs $\left(\beta_g^{\text{GA}}, \beta_g^{(0\rightarrow 7)}\right)$ out of all 16,172 age-related CpG sites that had the same sign, under the null hypothesis that the effects due to gestational age and chronological age were generated independently (see the Results section).

Assume the variance model for the data at birth and age seven is given by (1c) and let $r_g = \frac{\delta_g^2}{\delta_g^2 + \sigma_g^2}$. Then using the estimates for $\boldsymbol{C}_0, \boldsymbol{C}_{\text{diff}}$, along with the observed nuisance covariates $\boldsymbol{Z}_0, \boldsymbol{Z}_{\text{diff}}$,

$$\hat{\text{Corr}}\left(\hat{\beta}_g^{\text{GA}}, \hat{\beta}_g^{(0\rightarrow 7)}\right) = 0.66\left(1 - \hat{r}_g\right).$$

We then have that conditional on the true effects $\left(\beta_g^{\mathrm{GA}}, \beta_g^{(0\to7)}\right)^T$,

$$\left(\hat{\beta}_g^{\mathrm{GA}}, \hat{\beta}_g^{(0\to7)}\right)^T \approx N_2\left\{\left(\beta_g^{\mathrm{GA}}, \beta_g^{(0\to7)}\right)^T, \mathrm{diag}\left(c_g, d_g\right)\begin{pmatrix} 1 & 0.66\left(1-\hat{r}_g\right) \\ 0.66\left(1-\hat{r}_g\right) & 1 \end{pmatrix}\mathrm{diag}\left(c_g, d_g\right)\right\} \tag{S2}$$

for each $g \in [p]$, where $c_g, d_g$ are positive constants. Let $A_g$ be the event that CpG $g$ is an age-CpG at a 5% FDR. We assume that $A_g = \left\{|z|_g \geq t\right\}$, where $z_g = \hat{\beta}_g^{0\to7}/d_g$ is the z-score corresponding to $\hat{\beta}_g^{0\to7}$ and $t$ can be estimated as the smallest z-score with a q-value less than 0.05. The empirical distributions of $\left\{\hat{\beta}_g^{(0\to7)}\right\}_{g\in\{5\%\text{ FDR age CpGs}\}}$ and $\left\{\hat{\beta}_g^{GA}\right\}_{g\in\{5\%\text{ FDR gestational age CpGs}\}}$ were approximately symmetric around 0, which we took to imply $\left\{\beta_g^{(0\to7)}\right\}_{g\in[p]}$ and $\left\{\beta_g^{GA}\right\}_{g\in[p]}$ were symmetric around 0. For simplicity, we assume for density functions

$$h_{GA}(\cdot) = \sum_{r=1}^{R}\pi_r^{(GA)}N_1\left(\cdot; 0, \phi_r^{(GA)}\right) \qquad h_{(0\to7)}(\cdot) = \sum_{j=1}^{J}\pi_j^{(0\to7)}N_1\left(\cdot; 0, \phi_j^{(0\to7)}\right),$$

$\beta_g^{\mathrm{GA}} \overset{i.i.d}{\sim} h_{GA}(\cdot)$ and $\beta_g^{(0\to7)} \overset{i.i.d}{\sim} h_{(0\to7)}(\cdot)$. Such mixture normal densities can approximate a large class of parametric and non-parametric distributions [6]. Define $X_g, Y_g \in \mathbb{R}$ to be such that

$$\left(X_g, Y_g\right)^T \sim N_2\left\{0, \begin{pmatrix} 1 & 0.66\left(1-\hat{r}_g\right) \\ 0.66\left(1-\hat{r}_g\right) & 1 \end{pmatrix}\right\}.$$

Then under the null hypothesis that $\beta_g^{(0\to7)}$ and $\beta_g^{\mathrm{GA}}$ are independent and assuming (S2) is correct,

$$P\left\{\hat{\beta}_g^{\mathrm{GA}}\hat{\beta}_g^{(0\to7)} > 0 \mid A_g\right\} \leq \frac{P\left(\left\{X_g Y_g > 0\right\} \cap \left\{\left|Y_g\right| \geq t\right\}\right)}{P\left(\left|Y_g\right| \geq t\right)}.$$

We can easily estimate the above upper bound. Therefore, conditional on knowing whether or not each CpG is an age-associated CpG,

$$\mu = \mathbb{E}\left\{\sum_{g\in\{5\%\text{ FDR age CpGs}\}} 1\left(\hat{\beta}_g^{\mathrm{GA}}\hat{\beta}_g^{(0\to7)} > 0\right)\right\} = \sum_{g\in\{5\%\text{ FDR age CpGs}\}} P\left\{\hat{\beta}_g^{\mathrm{GA}}\hat{\beta}_g^{(0\to7)} > 0 \mid A_g\right\}$$

$\leq 14,236$

under the null hypothesis. Since the maximum variance for a Bernoulli random variable is $1/4$, an approximate lower bound for the test-statistic is

$$\frac{0.97 \times 16,172 - 14,236}{\sqrt{16,172/4}} = 23.3,$$

which has a corresponding $P$ value $\leq 10^{-119}$ under the normal approximation.

#### Determining the fraction of the variance in DNA methylation levels explained by maternal cotinine levels during pregnancy

Here we discuss our method for determining the fraction of the variance explained by maternal cotinine levels during pregnancy, which accounts for potential differences in the standard errors (i.e. sample sizes) of the maternal cotinine and genotype analyses.

The phenotype for maternal smoking was taken to be a factor variable with two levels, where the levels were smoker (cord blood plasma cotinine levels $\geq$ 10ng/mL) and non-smoker (cord blood plasma cotinine levels < 10ng/mL). The 10ng/mL threshold was chosen because it was the same cutoff used to define sustained maternal smoking in Joubert et al. [7]. We remark that 98% of the non-smoking mothers had cotinine levels below 2ng/mL, the limit of detection of the assay. We report results for DNAm levels at birth, although the results at age 7 are identical.

Let $c_i \in \{0, 1\}$ and $s_{gi} \in \{0, 1, 2\}$ be maternal smoking status and the genotype for the SNP closest to CpG $g$ for individual $i$, respectively. We assume that DNAm levels at birth at CpG $g$ in individual $i$ ($y_{gi}$) could be modeled as

$$y_{gi} = \mu_g + \beta_g^{(s)} s_{gi} + \beta_g^{(c)} c_i + \gamma_g^T \boldsymbol{z}_i + \epsilon_{gi}, \;\; \epsilon_{gi} \sim N\left(0, \sigma_g^2\right), \quad g = 1, \ldots, p; i = 1, \ldots, n, \tag{S3}$$

where $\boldsymbol{z}_i$ contain the nuisance covariates inferred genetic ancestry, sex, gestational age and methylation plate number. Using this model, we defined the fraction of the variance in DNAm levels at

CpG $g$ explainable by maternal smoking and genotype to be

$$\pi_g^{(c)} = \frac{\left\{\beta_g^{(c)}\right\}^2 \operatorname{Var}(c_i)}{\operatorname{Var}\left(y_{gi}\right)}, \quad \pi_g^{(s)} = \frac{\left\{\beta_g^{(s)}\right\}^2 \operatorname{Var}\left(s_{gi}\right)}{\operatorname{Var}\left(y_{gi}\right)}, \tag{S4}$$

respectively. We set $\operatorname{Var}(c_i) = p^{(c)}\left\{1 - p^{(c)}\right\}$ and $\operatorname{Var}\left(s_{gi}\right) = 2p_g^{(s)}\left\{1 - p_g^{(s)}\right\}$, where $p^{(c)}$ is the fraction of maternal smokers in our dataset and $p_g^{(s)}$ is the minor allele frequency for the SNP adjacent to CpG $g$. We remark that $p^{(c)} = 0.17$ was 2.8 times the national smoking during pregnancy (SDP) rate for non-Hispanic black mothers, 1.7 times the national SDP rate for non-Hispanic white mothers and 10.0 times the national SDP rate for Hispanic mothers [8], indicating $\pi_c^{(g)}$, depending on the exact population of interest, is likely an overestimate for the fraction of variance explained by maternal smoking. Since our goal was to determine the relative proportion of the variance explained by maternal smoking and genotype, i.e. $\pi_g^{(c)} / \left\{\pi_g^{(c)} + \pi_g^{(s)}\right\}$ and $\pi_g^{(s)} / \left\{\pi_g^{(c)} + \pi_g^{(s)}\right\}$, we need only estimate $\left\{\beta_g^{(c)}\right\}^2$ and $\left\{\beta_g^{(s)}\right\}^2$.

Since there was little detectable correlation between maternal smoking and the genotypes of SNPs adjacent to maternal smoking CpGs identified in Joubert et al. [7] (ms-CpGs), we ignored genotype when estimating $\beta_g^{(c)}$ and used McKennan et al. [5] to determine $\hat{\beta}_g^{(c)}$, an estimate for $\beta_g^{(c)}$, and subsequently used the method proposed in Stephens et al. [9] with only ms-CpGs as input to determine $\mathbb{E}\left[\left\{\beta_g^{(c)}\right\}^2 \mid \hat{\beta}_g^{(c)}\right]$, our estimate for $\left\{\beta_g^{(c)}\right\}^2$. To estimate $\left\{\beta_g^{(s)}\right\}^2$, we first computed $\hat{\beta}_g^{(s)}$, the ordinary least squares estimate for $\beta_g^{(s)}$ in Model (S3), using a random subset of 56% of the self-reported black children, and subsequently used Stephens et al. [9] with only ms-CpGs as input to determine $\mathbb{E}\left[\left\{\beta_g^{(s)}\right\}^2 \mid \hat{\beta}_g^{(s)}\right]$, our estimate for $\left\{\beta_g^{(s)}\right\}^2$. We used self-reported black children to avoid heterogeneous genetic effect sizes (due to population stratification), and only used 56% of those samples to determine $\hat{\beta}_g^{(s)}$ to ensure that the standard errors of $\hat{\beta}_g^{(c)}$ and $\hat{\beta}_g^{(s)}$ were approximately the same (Figure S5). This sub-sampling helped guarantee that the precision of the estimates $\mathbb{E}\left[\left\{\beta_g^{(c)}\right\}^2 \mid \hat{\beta}_g^{(c)}\right]$ and $\mathbb{E}\left[\left\{\beta_g^{(s)}\right\}^2 \mid \hat{\beta}_g^{(s)}\right]$ was the same, meaning that any difference in those estimates could not be attributed to the relatively small number of smokers in our study. We lastly plugged-in $\mathbb{E}\left[\left\{\beta_g^{(c)}\right\}^2 \mid \hat{\beta}_g^{(c)}\right]$ and $\mathbb{E}\left[\left\{\beta_g^{(s)}\right\}^2 \mid \hat{\beta}_g^{(s)}\right]$ for $\left\{\beta_g^{(c)}\right\}^2$ and $\left\{\beta_g^{(s)}\right\}^2$ into (S4) to estimate $\pi_g^{(c)} / \left\{\pi_g^{(c)} + \pi_g^{(s)}\right\}$ and

$\pi_g^{(s)} / \left\{\pi_g^{(c)} + \pi_g^{(s)}\right\}$.

#### Supplemental figures

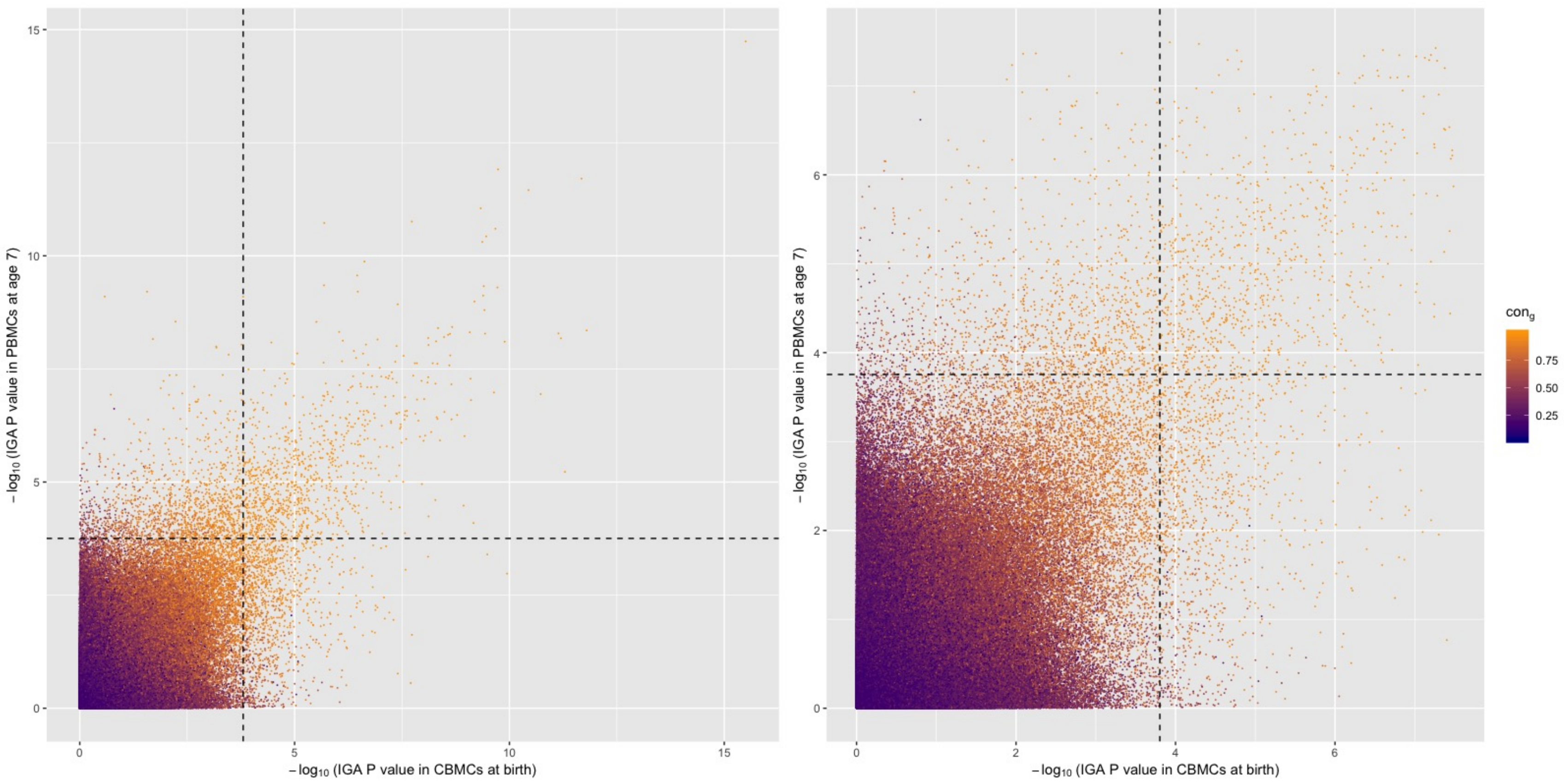


**Figure S1:** Relationship between inferred genetic ancestry (IGA) *P* values at birth and age 7 and estimated conserved sign rates ($con_g$). The *P* values were estimated using a standard regression model and the dashed lines indicate the 5% FDR threshold. The plot on the right is a zoomed-in version of the plot on the left.

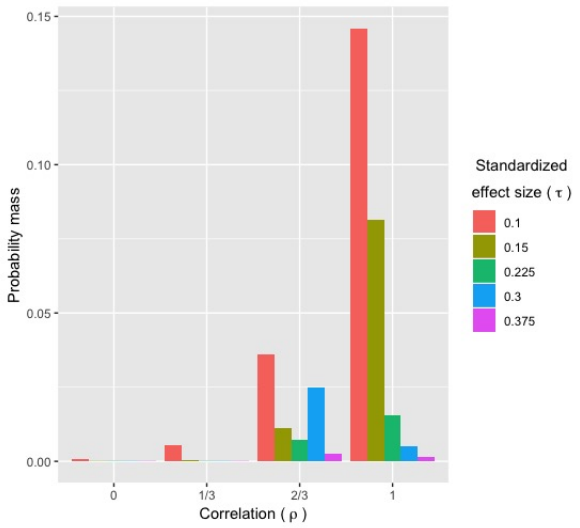


**Figure S2:** Probability mass of the components of $\hat{\boldsymbol{\pi}}_{(1,1)}$, where $\rho$ is the correlation between the reported race effect at birth and age 7 and $\tau$ is proportional to the expected magnitude of the effect sizes (see Model (1)). This trend was echoed in the inferred genetic ancestry analysis.

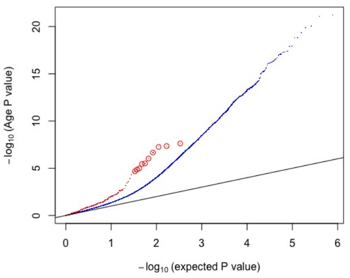


**(a)**

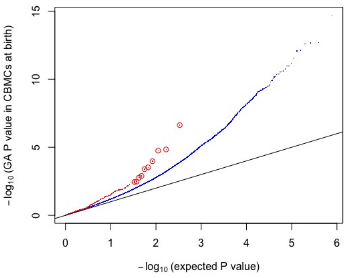


**(b)**

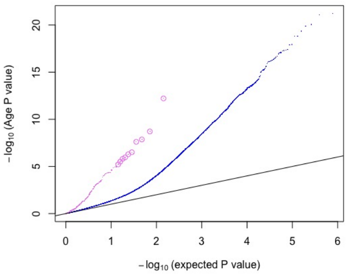


**(c)**

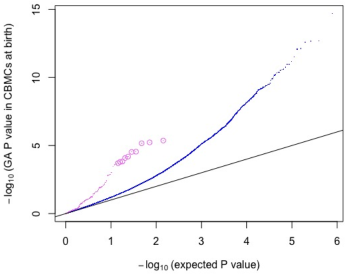


**(d)**

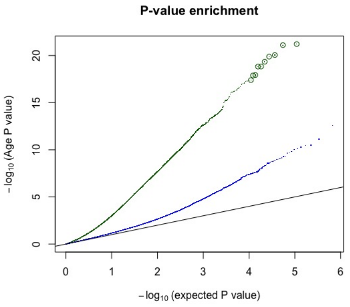


**(e)**

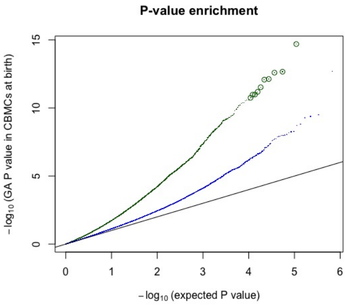


**(f)**

**Figure S3:** Distribution of *P* values for age (birth to age 7) (panels (a), (c) and (e)) and gestational age (GA) (panels (b), (d) and (f)). The red, violet and dark green dots in the upper, middle and lower panels are the 353 CpGs used to build the linear predictor of age in Horvath [10], the 148 CpGs used to build the linear predictor of gestational age in Knight et al. [11] and the 109,597 age (birth to age 5) CpGs discovered in Pérez et al. [12]. The blue dots are all of the other CpGs and the 10 enlarged circles are for visual aid.

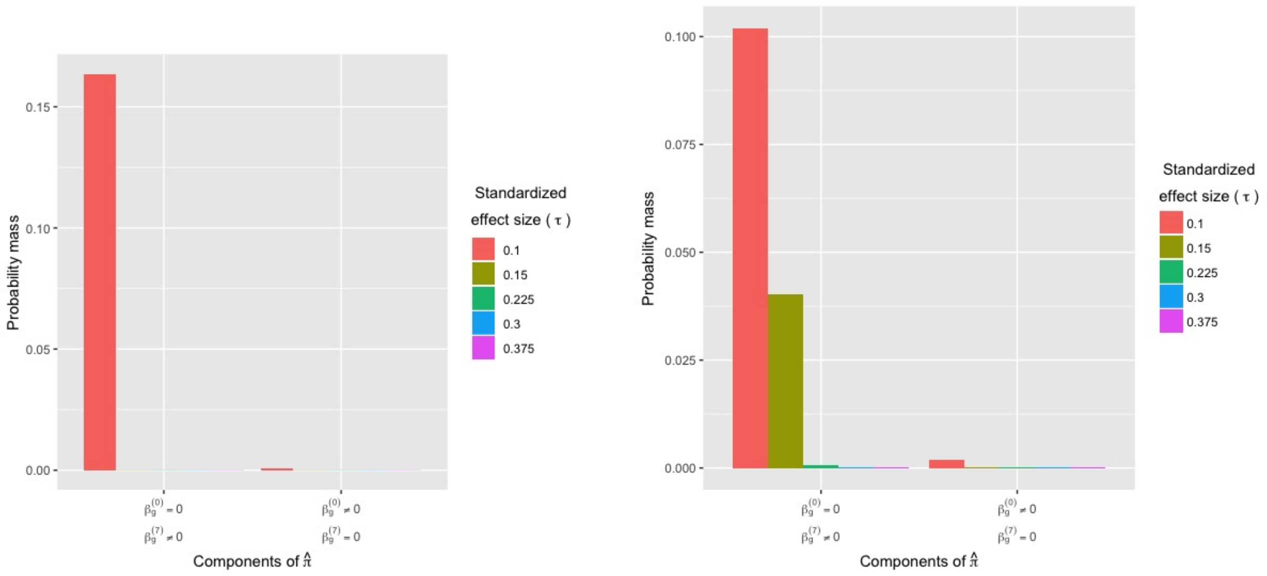


**(a)** **(b)**

**Figure S4:** Probability mass of components of $\hat{\pi}_{(1,0)}$ ($\beta_g^{(0)} \neq 0, \beta_g^{(7)} = 0$) and $\hat{\pi}_{(0,1)}$ ($\beta_g^{(0)} = 0, \beta_g^{(7)} \neq 0$) in the inferred genetic ancestry (a) and reported race (b) analyses.

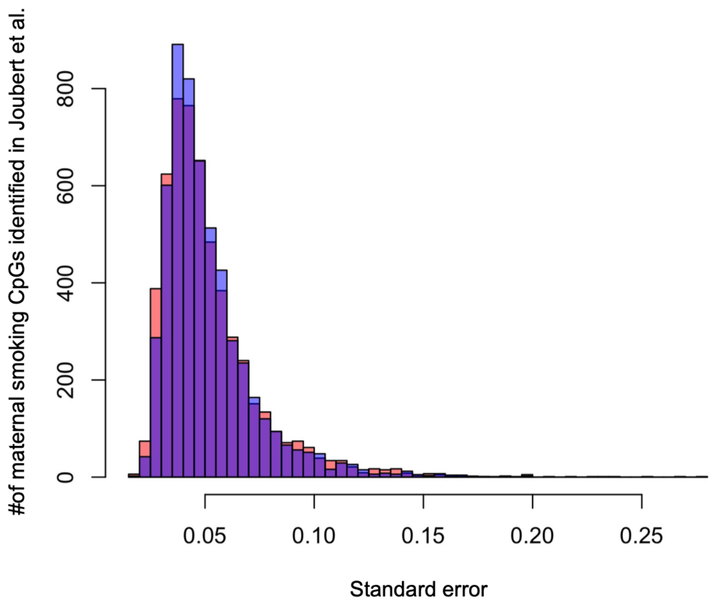


**Figure S5:** Histograms of standard errors for $\hat{\beta}_g^{(c)}$ (blue) and $\hat{\beta}_g^{(s)}$ (red) for all CpGs $g = 1, \ldots, 784, 484$ that are also maternal smoking-associated CpGs identified in Joubert et al. [7].
